# Decomposed frontal corticostriatal ensemble activity changes across trials, revealing distinct features relevant to outcome-based decision making

**DOI:** 10.1101/2024.03.23.586395

**Authors:** Takashi Handa, Tomoki Fukai, Tomoki Kurikawa

## Abstract

The frontal cortex-striatum circuit plays a pivotal role in adaptive goal-directed behaviours. However, the mediation of decision-related signals through cross-regional transmission between the medial frontal cortex and the striatum by neuronal ensembles remains unclear. We analysed neuronal ensemble activity obtained through simultaneous multiunit recordings in the secondary motor cortex (M2) and dorsal striatum (DS) while the rats performed an outcome-based choice task. Tensor component analysis (TCA), an unsupervised dimensionality reduction approach at the single-trial level, was adopted for concatenated ensembles of M2 and DS neurons. We identified distinct three spatiotemporal neural dynamics (TCA components) at the single-trial level specific to task-relevant variables. Choice-position selective neural dynamics was correlated with the trial-to-trial fluctuation of behavioural variables. This analytical approach unveiled choice-pattern selective neural dynamics distinguishing whether the incoming choice was a repetition or switch from the previous choice. Other neural dynamics was selective to outcome. Choice-pattern selective within-trial activity increased before response choice, whereas outcome selective within-trial activity increased following response. These results suggest that the concatenated ensembles of M2 and DS process distinct features of decision-related signals at various points in time. The M2 and DS may collaboratively monitor action outcomes and determine the subsequent choice, whether to repeat or switch, for coordinated action selection.

## Introduction

Animals can select an appropriate action based on sensory cues and outcomes of past actions to adapt flexibly to changing circumstances (Dolan & Dayan, 2013). The underlying neural circuit is widely believed to involve the frontal cortex-basal ganglia circuit (Ragozzino, 2007; Balleine & O’Doherty, 2010; Hikosaka & Isoda, 2010). However, it remains unclear how population neuronal dynamics interact between the frontal cortex and basal ganglia for the adaptive choice behaviour. Synchronous neuronal activity across the frontal cortex and downstream subcortical striatum is correlated with skilled motor learning (Koralek *et al*., 2013; Lemke *et al*., 2019) and flexible behaviours based on different rules in a T-maze (Oberto *et al*., 2022), as well as outcomes of action (Handa *et al*., 2021). Neural trajectories of the secondary motor cortex (M2), a part of the medial frontal cortex, and dorsal striatum (DS) concomitantly represent choice and outcome information during an outcome-based two-alternative choice task. Precise spike synchrony between M2 and DS neurons becomes more prominent during periods of improved task performance (Handa *et al*., 2021), suggesting cross-regional coactivation of neuronal population correlated with behavioural variables. However, little is known about how cross-regional neuronal dynamics contribute to decision-making during adaptive outcome-based action selection.

Large-scale neuronal activity can be analysed by adopting dimensionality reduction methods to profile the neuronal ensemble activity (Cunningham & Yu, 2014). Furthermore, conducting trial-by-trial analyses of ensemble activity provides a means to assess the relationship between alterations in the physiological aspects of neuronal ensemble activity and emerging behavioural variables within a single session (Quian Quiroga & Panzeri, 2009; Musall *et al*., 2019; Veuthey *et al*., 2020). Motivational states of an animal regulate behaviour and such internal states in the brain can be altered over trials within a single session (Burton *et al*., 1976; Berridge, 2004; Allen *et al*., 2019). For example, in the initial trials, a thirsty animal is notably motivated to engage in a behavioural task to attain a drop of water as a reward, whereas its motivated performance may diminish in the later trials.

Neural representation in the brain can be altered in accordance with such behavioural changes (Allen *et al*., 2019). We wonder whether the M2-DS ensemble adaptively processes decision-related information in correlation with behavioural variables during a single behavioural session.

To address these questions, we carried out simultaneous electrophysiological recordings in the M2 and DS while rats performed an outcome-based two alternative choice task. We analysed cross-region ensemble spike activity at the single-trial level by applying tensor component analysis (TCA) to the concatenated ensemble activity of M2 and DS neurons. TCA provides a crucial advantage over dimensional reduction approaches, allowing us to quantify trial-to-trial variations in neural activity. Additionally, because TCA is an unsupervised method, unexpected features of ensemble activity can be unveiled (Williams *et al*., 2018). We observed that choice-position selective neural dynamics were altered over trials, and such trial-by-trial alterations were correlated with trial-basis behavioural fluctuations. Certain neural dynamics revealed differential states between choice patterns, repetitive choices and switch choices, even when the incoming motor responses and outcomes were the same. Other neural dynamics could discriminate between the rewarded and unrewarded outcomes following action selection. Choice-pattern selective within-trial activity differed temporally from outcome-selective within-trial activity. These results suggest that trial-basis fluctuations in M2-DS ensembles could be attributed to behavioural variables and that the M2-DS ensemble was implemented in outcome monitoring and continued decision-making for action selection.

## Methods

### Animal preparation

All experiments were conducted in compliance with the Animal Experiment Plan of the Animal Experiment Committee of RIKEN (approval number: H25-2-234[1]). The multiunit recording results during task performance presented in this study were obtained from the reanalysis of previously collected behavioural and electrophysiological data (Handa *et al*., 2021). A brief overview of the experimental procedure for data acquisition related to this reanalysis is provided here, with comprehensive details available in the previous publication. Male Long-Evans rats (N = 14, 6 weeks, 200-220 g, Japan SLC, Inc., Japan) were employed. Home cages were situated in a temperature- and humidity-controlled environment with lights maintained on a 12-h light/dark cycle.

### Stereotaxic surgery

All surgical procedures were performed under sterile conditions. Rats were anaesthetised with 2% isoflurane, and their body temperature was monitored with a rectal probe and maintained at 37°C on a heating pad during the surgery. A sliding head attachment (Narishige, Japan) was implanted in the skull using a dual-curing resin cement (Panavia, Kuraray Noritake Dental) and dental resin (Unifast II, GC, Japan). Reference and grounding electrodes (Teflon-coated silver wire, A-M systems, U.S.A.) were positioned over the dura mater above the cerebellum. Following recovery from surgery, rats were deprived of water in their home cages, with water used as a reward for behavioural task execution; however, food was available *ad libitum*. Weekly, 10 ml of water was provided. To confirm recording sites within regions showing corticostriatal projections from the M2 to DS, a retrograde tracer, Fluoro-Gold (FG) (Fluorochrome), was injected into the DS 3 days before the electrophysiological recording experiment. A glass micropipette filled with 2% FG dissolved in 0.1 M cacodylic acid was installed on a micromanipulator angled medially by 27°. The pipette was inserted through a small burr hole drilled in the skull over left hemisphere (+1.5 mm to Bregma, 1.0 mm lateral to midline, 4.3 mm traveling distance). The pipette tip reached the dorso-central part of striatum (+1.5 mm anterior to Bregma, approximately 3.0 mm lateral to midline, approximately 3.8 mm ventral to pia mater) based on previous anatomical evidence (Reep *et al*., 2003). FG was iontophoretically infused using an iontophoresis pump (BAB-501; Kation Scientific, U.S.A). After the completion of training sessions, two cranial windows (1.2 mm diameter) were created above the DS and M2 of left hemisphere (+1.0 and +3.0 mm to Bregma, 3.0 and 1.0 mm to midline for DS and M2, respectively), and their dura maters were removed for electrophysiological recordings.

### Behavioural task

Rats underwent training to perform an outcome-based two-choice task using a customised multiple-rat training system (O’Hara &Co., Japan), facilitating parallel learning of the task paradigm for multiple rats simultaneously, as previously detailed (Handa *et al*., 2021). The behavioural task was controlled using custom-written software in LabVIEW (National Instruments, U.S.A.). Individual rats were secured in a body-supporting cylinder, and their heads were rigidly and painlessly fixed using a sliding head holder on a stereotaxic frame (Fig. 1A). Spouts were linked to a syringe on a single-syringe pump (AL-1000; World Precision Instruments, U.S.A.) using silicon tubing. Water delivery from each spout was regulated by a pinch valve and syringe pump triggered by the TTL signal. The trial initiated with a pure tone presentation (3 kHz, 1 s, 60-dB SPL) (‘Start’ in Fig. 1A). Rats were instructed to refrain from licking any spouts from the start cue until the appearance of another auditory cue (10 kHz, 0.2 s, 60-dB SPL) (‘Go’). If rats licked any spouts during the delay period (‘1st Delay’), the trial was promptly aborted. The pseudorandom delay period ranged from 0.7–2.3 s. Following the onset of the Go cue, rats could lick either the left or right spouts within a response window (5 s). The first lick was considered as a choice response (‘Choice’). If the chosen spout location aligned with the ongoing reward location, 0.1% saccharin water was delivered as a reward after a pseudorandom delay period ranging between 0.3 and 0.7 s (‘2nd Delay’). The next trial commenced after an outcome period (4 s, ‘Outcome’). However, when rats chose a no-reward spout, they received no sensory feedback but had an additional time of 5 s after the outcome period. The subsequent trial began after a timeout. Once the accumulated total number of rewarded trials reached 10 within each block, the reward-associated spout position reversed without any feedback, such as sensory or physical differences in the task. Block reversal occurred after approximately 11–12 trials if the rats frequently repeated the rewarded choice. If the number of unrewarded choices increased within a block, block reversal occurred after many more trials. Therefore, rats could not anticipate block reversal without experiencing forthcoming trials.

**Figure 1.**
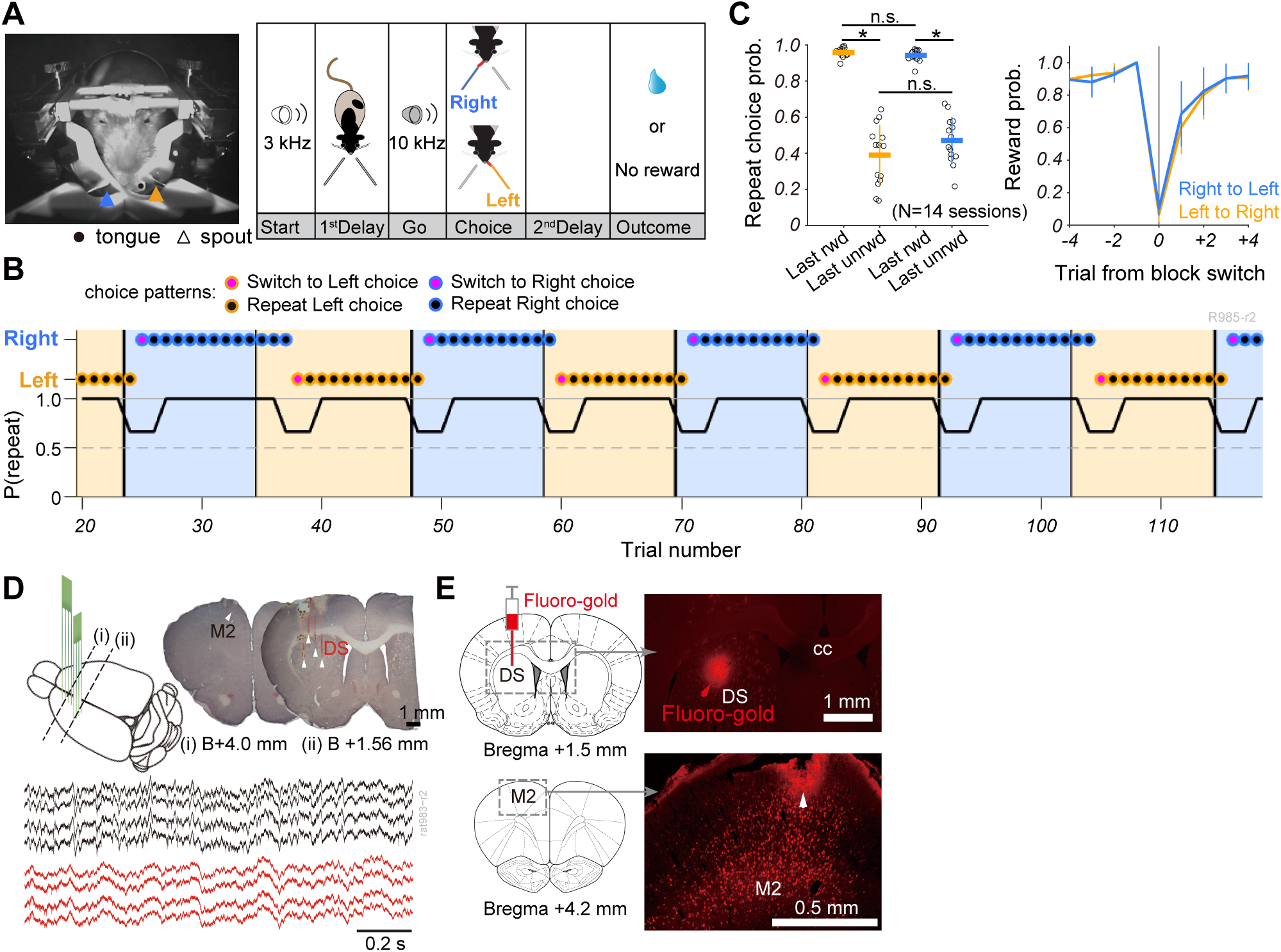
Flexible choice patterns revealing reward-guided repetitive choice and non-reward-guided switch choice**. (A)** Left: Snapshot of a head-fixed rat at the moment of licking choice toward a left spout (orange arrowhead). A black circle and blue arrowhead indicate the position of the tongue and right spout, respectively. Right: Schematic illustration of an outcome-based two alternative choice task. Each trial began with the presentation of an auditory cue (Start, 3 kHz). Rats awaited another auditory cue (Go, 10 kHz) while abstaining from licking during the initial delay period (1st Delay). Subsequently, they chose either left or right spouts by licking within 5 s. A reward was provided after the second delay period (0.3–0.7 s) if the selected spout-position aligned with the current reward-position; otherwise, no reward was given, accompanied by a lack of sensory feedback and a 5-s timeout. **(B)** Representative choice pattern revealing repetitive choice behaviour post-reward acquisition in last trial and switch choice behaviour post-unrewarded trials. Background colours indicate ongoing reward-positions (orange: left spout, blue: right spout). Thick and thin vertical lines denote reversals of reward-position: from left to right and from right to left, respectively. Colours of outline and face in symbols represent choice-position (left or right) and choice-pattern (repetitive or switch), respectively. A line plot displays trial series of the probability of repetitive choice (average probability over three trials). **(C)** Left: Probability of repetitive choice (orange: left choice, blue: right choice) after rewarded (Last rwd) and unrewarded (Last unrwd) outcomes in the last trial. Circles indicate individual sessions (14 sessions, 12 rats). Horizontal lines and error bars represent mean and SD, respectively. Statistical significance was confirmed by one-way ANOVA followed by *post hoc* Tukey-Kramer test (*: *P* <0.001). Right: Averaged choice patterns around the reversal of reward-position (orange: from left to right, blue: from right to left). Values present mean and SD. **(D)** Top: Schematic illustration of recording sites with two probes in M2 and DS of left hemisphere. Recording sites (white arrowhead) in M2 and DS in the Nissl-stained coronal brain sections. Bottom: Representative local field potentials simultaneously recorded from M2 (black) and DS (red). **(E)** *Post hoc* confirmation of injection site of retrograde tracer, fluoro-gold (red arrowhead), in DS and the corticostriatal projection neurons labelled with fluoro-gold in M2, including the recording site (white arrowhead). cc: corpus callosum. B (or Bregma) indicates AP coordinate based on the rat brain atlas (Paxinos and Watson, 2009).

### Electrophysiological recordings during task performance

Following 21 training sessions, each animal underwent two daily recording experiments. Multi-neuron activity was simultaneously recorded from the M2 and DS of the left hemisphere using two 32-channels silicon probes. These probes consisted of four shanks (0.4 mm shank separation), each featuring tetrode-like electrode sites spaced vertically by 0.5 mm (A4x2-tet-7/5 mm-500-400-312, NeuroNexus Technologies, U.S.A.). Each probe was connected to a custom-made headstage on one of two fine micromanipulators (1760-61; David Kopf Instruments, U.S.A.) mounted on a stereotaxic frame (SR-8N, Narishige, Japan). The silicon probe was vertically inserted (depth from pia mater: 1.2 mm) into M2 (at the centre of probe: +3.0–3.6 mm to Bregma, 1.0–1.4 mm to midline), with the shanks aligned along the midline (Fig. 1D). Another silicon probe, angled posteriorly by 6°, was inserted into DS through a cranial window (at the centre of probe: +0.6–1.0 mm to Bregma, 2.7–3.1 mm to midline, 4.0 mm traveling distance), with the shanks aligned along the coronal suture (Fig. 1D). Multiunit signals were amplified by the headstages before being fed into main amplifiers (Nihon Koden, Japan) with a band-pass filter (0.5 Hz–10 kHz). All neural data were sampled at 20 kHz using two hard-disc recorders (LX-120, TEAC, Japan), capturing the time of the task and licking events for each spout (left and right).

### Histology

After the recording sessions, rats were deeply anaesthetised with urethane (2–3 g/kg, i.p.) and subsequently perfused intracardially with chilled saline followed by 4% paraformaldehyde (PFA) dissolved in 0.1 M phosphate buffer (PB). The fixed brains were stored in 4% PFA overnight and then placed in a 30% sucrose solution in 0.1 M PB for 2 weeks. Postfixed brains were frozen and coronally sliced into 50-μm thick serial sections using a microtome Cryostat (HM500OM, Microm, U.S.A.). The brain sections were stored in 0.1 M PB at 4°C overnight. For fluorescent visualisation of FG-labelled neurons, brain sections were incubated with an anti-FG antibody from rabbit (AB153, 1:3000, Millipore) at 4°C overnight. Subsequently, they underwent incubation with goat anti-rabbit IgG conjugated with Alexa fluor-594 (A11012, 1:500, Invitrogen) for 2 h. Fluorescence images were acquired using a fluorescence microscope (Olympus, AX70, Japan) to confirm the presence of FG-labelled neurons around the silicon probe recording locations in the M2 and near the FG injection site in the DS. To verify the silicon probe track, the slices were counterstained with Neural red Nissl. Recording locations in the M2 and DS as well as AP coordinates were determined in accordance with the rat brain atlas (Paxinos and Watson, 2009).

### Data analysis

All behavioural and neuronal data were analysed by custom-written MATLAB scripts (The MathWorks Inc., U.S.A.).

### Data set

We reanalysed behavioural and multi-neuron spike datasets previously recorded, sorting them into 14 recording sessions with 12 rats using a new analytical approach. Details about spike sorting and clustering were described in the previous publication (Handa *et al*., 2021). The sessions are as follows, with session-ID, reward acquisition probability (*P*), and the number of well isolated cells in M2/DS: R982-r1 (*P* = 0.828, M2/DS = 45/43), R983-r1 (*P* = 0.837, M2/DS = 35/55), R983-r2 (*P* = 0.834, M2/DS = 58/48), R985-r1 (*P* = 0.853, M2/DS = 30/60), R985-r2 (*P* = 0.867, M2/DS = 66/26), R986-r1 (*P* = 0.813, M2/DS = 16/20), R991-r1 (*P* = 0.806, M2/DS = 46/31), R997-r1 (*P* =0.740, M2/DS = 24/34), R1000-r1 (*P* = 0.784, M2/DS = 12/32), R1002-r1 (*P* =0.744, M2/DS = 31/11), R1004-r1 (*P* = 0.792, M2/DS = 20/51), R1005-r1 (*P* = 0.794, M2/DS = 65/64), R1009-r1 (*P* = 0.774, M2/DS = 35/33), R1012-r1 (*P* = 0.755, M2/DS = 40/26).

### Tensor component analysis (TCA)

To generate an original neural data tensor ***X*** for each recording session, we computed trial-based peri-event time histograms aligned at the choice response, using a 200-ms sliding window with a 50-ms step for individual M2 and DS neurons. Subsequently, we obtained a third-order tensor (neuron, time, and trial) for the M2-DS ensemble through TCA (Fig. 2A). To investigate the single-trial dynamics of the M2-DS ensemble activity, we applied TCA to tensor ***X*** using a MATLAB-based toolbox (Tensor Toolbox for MATLAB, version 3.2.1, https://www.tensortoolbox.org/) (Bader & Kolda, 2008). In this analysis, the M2-DS ensemble activity was decomposed into a third-order tensor 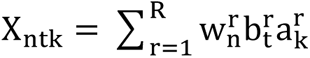 by the summation of one-rank components (Fig. 2A). Each component consists of three vectors: 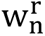 represents the n-th element of a ‘neuron factor’ vector, reflecting a prototypical firing rate pattern across neurons; 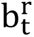 is t-th element of a ‘temporal factor’ vector, representing a temporal basis function across time; and 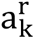 is the k-th element of a ‘trial factor’ vector, signifying a trial-specific bias for spatiotemporal activity in a trial. We set the number of TCA components R to 15 for canonical polyadic decomposition based on a previous study, which suggested that 15 components were sufficient to profile the population of neurons encoding task variables in a behavioural experiment (Williams *et al*., 2018). For the analysis of ensemble coding in M2- and DS-alone, we applied TCA to the M2 and DS ensembles alone, respectively.

**Figure 2.**
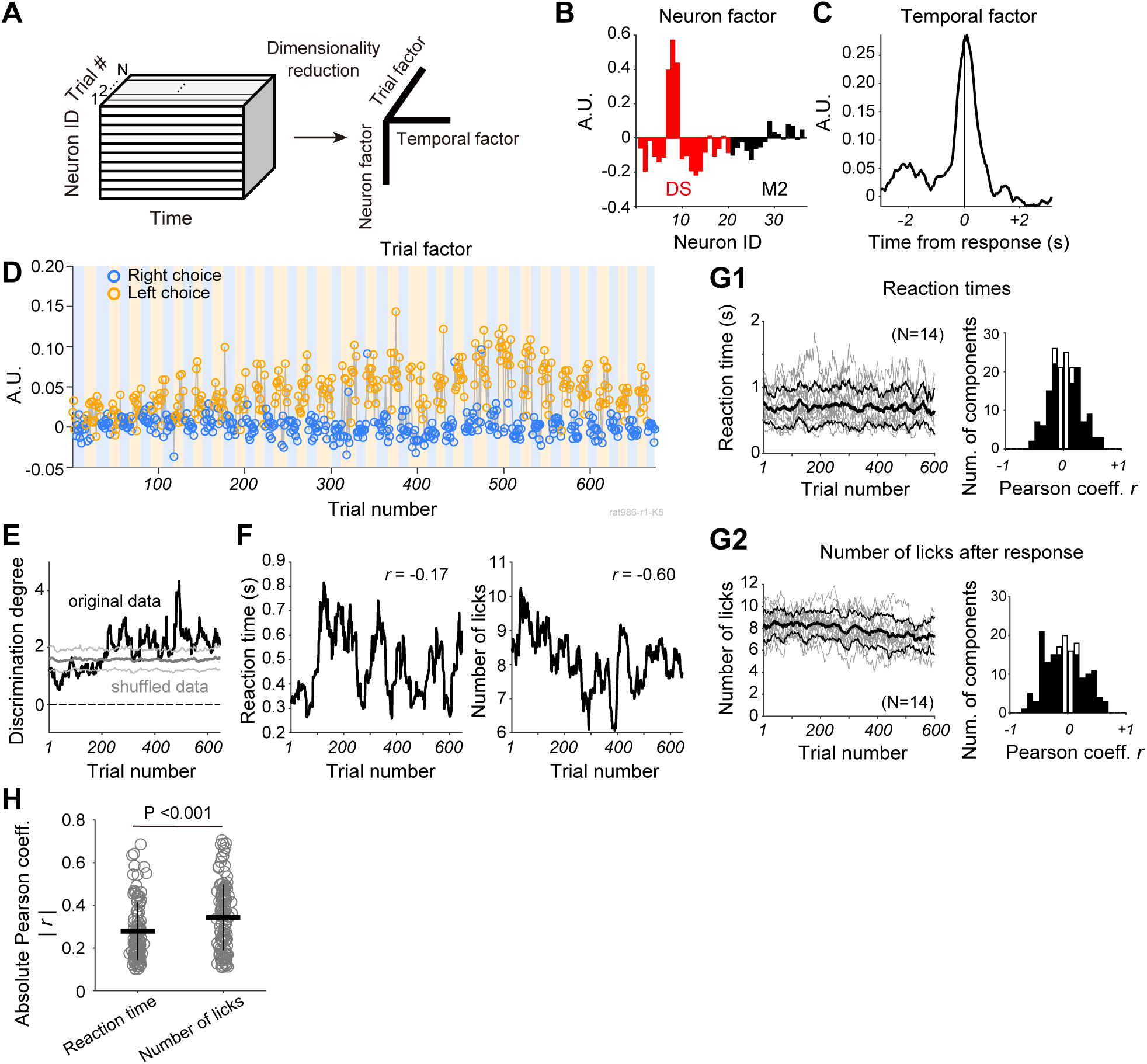
Trial-to-trial changes in choice-position selective TCA component of M2-DS ensemble activity are related to behavioural variables. **(A)** A schematic illustration of tensor component analysis (TCA). **(B–D)** Example of a TCA component. **B**: Neuron factor, including M2 (black) and DS (red) neurons. **C:** Temporal factor (within trial activity). **D**: Trial factor of the TCA component. Symbol colours denote the choice position at each trial (left: orange, right: blue). Background colours represent the ongoing reward-position (orange: left spout, blue: right spout). **(E)** Trial series of discrimination degree of the trial factor shown in D using the original trial order (black) and shuffled trial order (grey, mean and SD of discrimination degree acquired by repeating shuffling trial order). **(F)** Left: Trial series of reaction times (RTs), and (right) trial series of the number of licks after the choice response in the same session as shown in B–D. ‘*r’* denotes Pearson’s correlation coefficient. **(G1)** Left: Trial-series of RTs over 14 sessions. Grey and black lines represent individual and averaged values. Error range indicates SD. Right: The distribution of correlation coefficients between discrimination degree and RTs. **(G2)** Left: Trial-series of the number of licks after the choice response over 14 sessions. Right: The distribution of correlation coefficients between discrimination degree and the number of licks. Open and filled bars indicate the number of non-significant and significant correlation coefficients, respectively (Pearson’s correlation, *P* <0.01). **(H)** Comparison of absolute correlation coefficients showing statistical significance (as shown in G1 and G2) between two behavioural variables. Statistical significance was assessed using a two-sample t-test. Horizontal and vertical lines represent mean and SD, respectively.

### Choice-position selectivity while trials progress

We assessed significant differences in neural dynamics between left- and right-choice trials in a given session by applying a two-sample t-test (*P* <0.05) to trial factors for each TCA component. If the differences were statistically significant, we referred to the TCA component as ‘choice-position’ selective TCA component. We computed the discrimination degree *d’* as a choice-position selectivity index in a window of 30 trials by sliding the window with one trial step. Instantaneous *d’* was computed as follows:

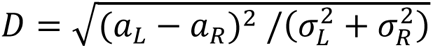

where *aL* and *aR* indicate the means of the trial factors of the left and right trials within 30 trials, respectively. *sL* and *sR* denote the standard deviations (SD) of the trial factors of the left and right trials within 30 trials, respectively.

If *a*L - *a*R >0, *d’* = +D

If *a*L - *a*R <0, *d’* = -D

To assess whether the trial-wise discrimination degree (original *d’*) was altered across trials, we used a permutation test by comparing it with the control data. As a control discrimination degree, we randomly shuffled the trial order and computed the trial-wise of discrimination degree (surrogated *d’*) by means of the shuffled trial factor. Subsequently, the original *d’* was compared with the surrogated *d’* by paired t-test (*P* <0.05). This procedure was repeated 100 times. If a statistically significant difference was observed in >95% of 100 repetitions, we determined that the TCA component altered the choice-position selectivity across trials.

### Correlation of choice-position selectivity with behavioural variables

To investigate whether the change in choice-position selectivity of the TCA component was correlated with the change in behavioural variables, we calculated two behavioural variables: reaction times (RTs) and the number of licks after the choice response. In each trial, RT was calculated as the duration between Go cue onset and the time at which the first lick occurred after Go cue onset, and the number of licks that emerged within 2 s after the choice response. We computed the average RTs and number of licks by sliding the analysis window of 30 trials in one trial step. We subsequently computed Pearson’s correlation between the trial series of choice-position selectivity *d’* and each behavioural variable. If the P-value was <0.01, we defined the TCA component as significantly correlated with the behavioural variable.

### Quantification of choice-pattern selectivity in M2-DS combined ensemble activity

To quantify the extent to which the neural dynamics in switch-choice trials differed from those in repetitive-choice trials, we computed the standard score (Z-score) of trial factors based on their mean and SD within individual blocks. In the case of the left block, during which the left spout was associated with reward delivery, we calculated the Z-score of trial factors in individual left blocks, ranging from the trial where the rat switched its choice from the preceding unrewarded choice (switching from right unrewarded choice to left rewarded choice) to the last trial before the reversal of reward position. The Z-scores in the switch-choice trial (Switch) and in the three continued repetitive-choice trials (Rep-1, Rep-2, and Rep-3) were computed as follows:

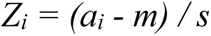

where *ai*, *m,* and *s* indicate a trial factor (*i* = Switch, Rep-1, Rep-2, or Rep-3 choice trial) in a block, and the mean and SD of the trial factors within the block, respectively. To derive *m* and *s* in individual blocks (for example, the left reward block), we used trials that included the same choice (left choice) and outcome (rewarded) conditions, excluding other choice conditions (right choice and unrewarded). In a given block, if any unrewarded choice trials intermingled before the four continued rewarded choices were achieved, the block was removed from this analysis. We assessed the statistical significance of differences in Z-scores among the four conditions (Switch, Rep-1, Rep-2, and Rep-3) by one-way ANOVA, followed by *post hoc* multiple comparison Dunnett’s test (*P* <0.05). If all of three pairs (Switch vs. Rep-1, Switch vs. Rep-2, and Switch vs. Rep-3) exhibited significance, we categorised the TCA component as a ‘choice-pattern’ selective TCA component. The same computation was performed separately for the right reward block.

### Quantification of outcome selectivity in M2-DS combined ensemble activity

To quantify the level of trial factor differences between rewarded and unrewarded choice trials in M2-DS combined ensemble activity, Z-scores of trial factors were computed, similar to the aforementioned analysis of choice-pattern selectivity. For the left block, trial factors were collected in a sequence of trials beginning from a trial where the rat correctly switched its choice from the preceding unrewarded choice (right to left) up to one trial before the initial switch-choice trial (left to right) in the next block. This chunk of trials encompassed both repetitive-choice but unrewarded trials, occurring due to the reversal of the reward position. Z-scores were computed for the initial three repetitive choice and reward trials (Rep-1 & rwd, Rep-2 & rwd, and Rep-3 & rwd), along with those of the repetitive choices and unrewarded trials (Rep & unrwd), as described earlier. If any unrewarded choice trials were interspersed before achieving the four consecutive rewarded choices, the block was excluded from the analysis. Assuming that Rep-1 and rwd, Rep-2 and rwd, and Rep-3 & rwd, and Rep and unrwd represented the same choice pattern (repetitive choice) but different outcome conditions (Rep-1 & rwd, Rep-2 & rwd, and Rep-3 & rwd: rewarded; Rep & unrwd: unrewarded). Statistical significance was determined through one-way ANOVA among the four choices, followed by *post hoc* multiple comparison Dunnett test (*P* <0.05). If all three pairs (Rep & unrwd vs. Rep-1 & rwd, Rep & unrwd vs. Rep-2 & rwd, and Rep & unrwd vs. Rep-3 & rwd) exhibited significance, the TCA component was categorised as ‘outcome’ selective TCA component. A parallel computation was conducted for right-choice trials separately.

### Quantification of choice-pattern and outcome selectivity in M2- and DS-alone ensemble activities

TCA was applied separately to the M2- and DS-alone ensemble activities using the same dataset as the M2-DS ensemble activity. Z-scores were computed for the analyses of choice-pattern and outcome selectivity, following the previously described methods.

### Statistics

To examine repetitive choice probability after rewarded and unrewarded trials, repetitive choice probability data were assessed using one-way ANOVA, followed by *post hoc* Tukey-Kramer test (*P* <0.05). The choice-position selective TCA component was evaluated using a two-sample t-test (*P* <0.05). Significant differences in the degree of discrimination between original and trial-order shuffled data were determined using a paired t-test (*P* <0.05) with 100 repetitions, and statistical significance was declared if observed >95 times. Significant correlations between behavioural variables and the degree of discrimination were assessed using Pearson’s correlation analysis (*P* <0.01). Absolute correlation coefficients were compared using a two-sample t-test (*P* <0.05). Choice-patten related TCA and outcome-related TCA were determined using one-way ANOVA (*P* <0.05), followed by *post hoc* Dunnett’s test (*P* <0.05). The peak time of the temporal factor was compared between choice-pattern related and outcome-related TCA components using the Mann-Whitney U test (*P* <0.05). Mean absolute Z-scores were compared among the M2-DS ensemble-based TCA data, M2 ensemble-based TCA data, and DS ensemble-based TCA data using one-way ANOVA (*P* <0.05). P-values considered statistically significant are indicated in parentheses for each statistical test described earlier.

## Results

### Rats exhibit retrospective outcome-based choices

The findings present a re-analysis of previously published behavioural and electrophysiological data (Handa *et al*., 2021) using a novel analytical approach. Head-restrained rats engaged in an outcome-based two-choice task, selecting between two spouts to make their choice (Fig. 1A). A reward was received when the chosen spout matched the ongoing reward-spout position. The action-outcome association was systematically reversed without sensory feedback after accumulating >10 rewarded trials in each block. We analysed the behavioural data from 14 recording sessions with 12 rats. Across all sessions, the average number of trials per session was 589 ± 124 (mean ± SD). Rats discerned the reversal of choice-reward contingency within subsequent trials, responding to the experience of no-reward events, time-out, and reward acquisition (Fig. 1B). In both the left and right choice trials, rats repeatedly selected the same spout as that in the preceding rewarded trial but switched their choice following one to several unrewarded trials (Fig. 1B and C) (one-way ANOVA: *P* <10^-21^, *post hoc* Tukey-Kramer test: last rewarded trials vs. last unrewarded trials, left choice, *P* <0.001, right choice, *P* <0.001). No statistically significant differences were observed in the repetitive choice probabilities between the left- and right-choice trials regardless of the outcome conditions in the preceding trial (*post hoc* Tukey-Kramer test: last rewarded trials, left choice vs. right choice, *P* = 0.973; last unrewarded trials, left choice vs. right choice, *P* = 0.201). The rats demonstrated an inability to predict the reversal of the reward block, choosing nearly no reward-associated spouts in some trials post-reversal. In response to the block reversal, rats then retrospectively switched to another choice (Fig. 1C, right). In essence, the choice pattern of the rats closely mirrored the win-stay and lose-shift strategy, an optimal approach for maximising reward acquisition in the current task, considering that the rats did not anticipate the block reversal.

### Simultaneous recording of ensemble neural activity from M2 and DS

To investigate whether frontal corticostriatal ensembles are encoded at the single-trial level, we conducted simultaneous recordings of multi-neuron activity in both the M2 and DS of the left hemisphere using two multi-electrode probes (Fig. 1D). Across 14 recording sessions, a considerable number of units were recorded in both M2 (mean ± SD, 37.3 ± 16.5 units) and DS (38.8 ± 15.5 units). The rodent M2 (or the rostral agranular medial cortex) is one of the primary cortical areas projecting to the dorso-central region of the striatum (Reep *et al*., 2003; Cheatwood *et al*., 2003; Hintiryan *et al*., 2016). To confirm the presence of such corticostriatal projections from the recording site in M2 to that in the DS, we iontophoretically infused a retrograde tracer Fluoro-Gold (FG) into the central part of the DS before recording sessions (Methods). FG-labelled corticostriatal neurons were primarily observed in layers 3 and 5 of the M2 (Fig. 1E). The probe track for M2 and DS recordings aligned with the FG-labelled region in the M2 or near the injection site in the DS (Fig. 1D and E), confirming the recording of the multi-neuron activity from the directly connected subregions of the M2 and DS.

### Trial-by-trial changes in choice-position selective activity of M2-DS ensembles are correlated with behavioural variables

To investigate the dynamic changes in the characteristics of M2-DS ensembles across trials, we utilised TCA (Williams *et al*., 2018)) for both dimensionality reduction of high-dimensional neuronal activity and quantification of trial-to-trial fluctuations in ensemble activity (Methods). Here, we refer to the collective firing activity of multiple M2 and DS neurons as the M2-DS ensemble. TCA decomposes the M2-DS ensemble activity into a third order tensor 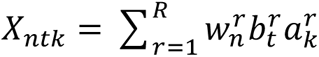 by the summation of one-rank component (Fig. 2A). Each component comprises three vectors: 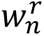 is the n-th element of a ‘neuron factor’ vector, representing a prototypical firing rate pattern across neurons (Fig. 2B); 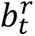 is the t-th element of a ‘temporal factor’ vector, indicating a temporal basis function across time (Fig. 2C); and 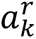 is the k-th element of a ‘trial factor’ vector, serving as a trial-specific bias for spatiotemporal activity in a trial (Fig. 2D).

We hypothesised that the TCA components of the M2-DS ensemble would exhibit choice-position selectivity that changes across trials, consistent with our previous findings of choice-position selectivity in single-unit and population activity in both the M2 and DS during this task (Handa *et al*., 2021). Indeed, the trial factor of this TCA component revealed significant differences between the left and right choices (two-sample t-test, *P* <10^-126^). Additionally, we observed that the difference in trial factor between choice positions was small in the initial trials but gradually increased in the middle and later parts of this session (Fig. 2D). To quantify the gradual changes in trial factors across trials, we calculated the degree of discrimination across trials as choice-position selectivity (Methods). The discrimination degree increased across trials and significantly differed from the discrimination degree derived by shuffling trial order (Permutation test, *P* <0.01; Methods) (Fig. 2E).

We examined whether the gradual changes in the discrimination degree were associated with alterations in behavioural variables across trials. Specifically, we investigated the relationship between the discrimination degree and two behavioural variables: reaction time (RTs) and the number of licks after the choice response. The trial series of the number of licks exhibited a higher correlation with the degree of discrimination of the TCA component (Pearson’s correlation: *r* = -0.60, *P* <0.0001) than with the trial series of RTs (*r* = -0.17, *P* <0.0001) (Fig. 2F). Although RTs and the number of licks varied across the 14 sessions, an average reduction in the number of licks was observed across trials (Fig. 2G2). The behavioural results suggest that this reduction in the number of licks over the trials may reflect changes in the motivational state of the rats to engage in task performance. Among the 166 choice-position selective TCA components, 112 (67.4%) and 125 (75.3%) were significantly correlated with RTs and the number of licks, respectively (Pearson’s correlation, *P* <0.01). Among these significantly correlated TCA components, 87 revealed a significant correlation with both RTs and the number of licks. The magnitudes of the significant correlation coefficients were larger in the correlation with the number of licks than in the correlation with RTs (two-sample t-test, *P* <0.001) (Fig. 2H). These results indicate that TCA unveils the dynamics of choice-position selectivity in M2-DS ensembles across trials, which may be linked to changes in behavioural variables such as motor preparation and/or motivational state.

### TCA unveils activity patterns of M2-DS ensembles distinguishing between repetitive and switch choices

In this study, TCA not only confirmed the anticipated choice-position selective activity type of M2-DS ensembles, as revealed in our previous analysis (Handa *et al*., 2021), but also unveiled an unexpected trial factor pattern that differentially altered depending on the choice pattern—switch choice versus repetitive choice (Fig. 3A-C). A representative TCA component exhibited an increase and decrease in within-trial activity (temporal factor) before and after the response, respectively (Fig. 3B). The trial factor displayed distinct patterns between repetitive and switch choices in right-choice trials but not in left-choice trials (Fig. 3C). In right-choice trials, the trial factors in switch trials significantly deviated from those in the repetitive trials (Fig. 3C and D). In contrast, in left-choice trials, the trial factor fluctuated similarly in both switch and repetitive choices. To quantify the deviation within a single block of trials, we calculated the Z-scores of trial factors within each block to compare switch choice, first, second, and third repetitive choices, where the movement direction and outcome condition (rewarded) were the same (Fig. 3E top, Methods). The Z-scores in switch-choice trials significantly differed from those in all other repetitive choice trials in the right (contralateral) choice condition (one-way ANOVA: *P* <10^-13^, *post hoc* Dunnett test, Switch vs. Rep-1: *P* <0.0001, Switch vs. Rep-2: *P* <0.0001, Switch vs. Rep-3: *P* <0.0001), whereas the Z-score was not significantly different in the left (ipsilateral) choice condition (one-way ANOVA: *P* = 0.102) (Fig. 3E). The lateralized difference between repetitive and switch choices suggests that choice-pattern selective neural dynamics did not account for the differences in the previous choices or outcomes between the choice patterns. Choice-pattern-selective TCA components were more frequently observed in right (contralateral) choice trials (N = 21) than in left (ipsilateral) choice trials (N = 13). Four TCA components displayed choice-pattern selective activity in both left and right choice trials. The right choice-preferred TCA component (11 sessions) was detected in more recording sessions than the left choice-preferred TCA component (six sessions) (Fig. 3F). The TCA component revealing choice-pattern selectivity in both left and right choices was observed in three sessions. The population of Z-scores for the choice-pattern-selective TCA trial factor was significantly higher in switch choice trials than in continued repetitive choice trials (Fig. 3G). This result suggests that the M2-DS ensembles differentially encode incoming choice information between switch choice (left to right) and repetitive choice (right to right) at the single-trial level.

**Figure 3.**
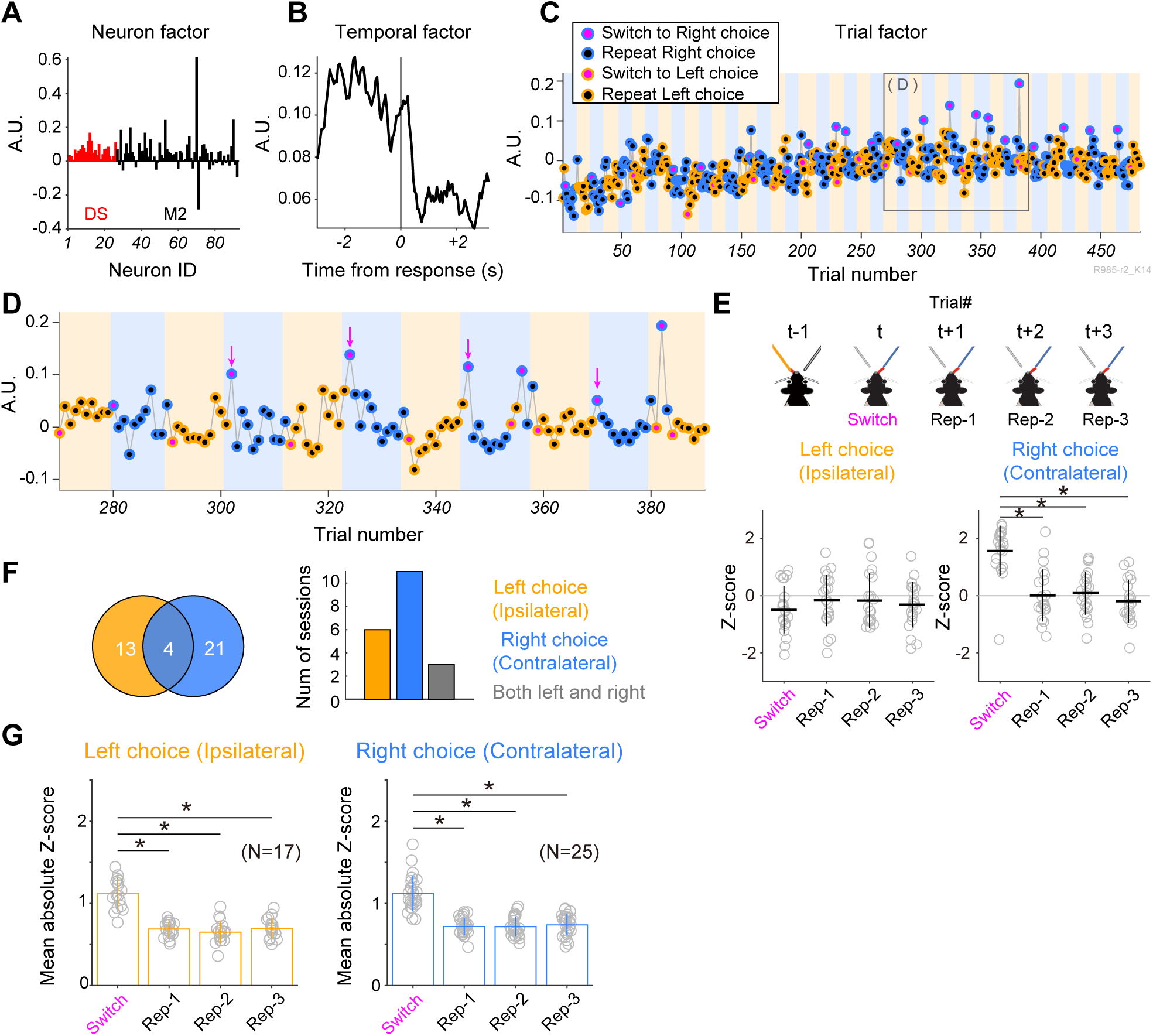
Trial-order activity pattern of M2-DS ensembles related to repetitive and switch choices. **(A)** Neuron factor, **(B)** temporal factor, and **(C)** trial factor of a representative TCA component revealing large variance in switch choice trials. Edge and face colours represent choice-positions (orange: left, blue: right) and choice-patterns (black: repetitive choice, magenta: switch choice), respectively. Background colours denote ongoing reward positions (orange: left spout, blue: right spout). A grey box presents an area enlarged in D. **(D)** Representatives of switch choice trials (magenta arrows) exhibit significantly larger variance when the ongoing reward position is at the right spout. **(E)** Top: A schematic illustration of choice-patterns utilized for the statistical estimation of differences in trial factors among repetitive and switch choices. This trial sequence illustrates a switch from left to right choices after the reversal of reward position (from the left spout to right the spout). At trial t, the animal shifts its choice to the right spout, transitioning from the left choice selected in the preceding trial (t-1). Subsequent repetitive choices (t+1, t+2, and t+3) are employed to compare differences among switch (Switch) and repetitive (Rep-1, Rep-2, and Rep-3) choice patterns. Bottom: Z-scores of trial factors in Switch, Rep-1, Rep-2, and Rep-3 choice patterns within the same dataset as depicted in C and D, corresponding to the left (ipsilateral) and right (contralateral) choices of the rat. The statistical significance of differences is assessed through one-way ANOVA followed by *post hoc* Dunnett test (*: *P* <0.05). Horizontal and vertical lines represent mean and SD, respectively. **(F)** The Venn diagram illustrates the number of TCA components with significantly different Z-scores of trial factors between repetitive choice and switch choice trials in left choice (orange), right choice (blue), and both (merge) conditions. The bar graph indicates the number of sessions featuring TCA components with significantly different Z-scores of trial factors between repetitive and switch choice trials. **(G)** Population data of TCA components reveal a significant difference in Z-scores between repetitive and switch choice trials. Individual dots indicate the mean of absolute Z-score per TCA component. Bar graphs show the mean and SD. Statistical significance of the difference is assessed through one-way ANOVA followed by *post hoc* Dunnett test (*: *P* <0.05).

### TCA component exhibits a differential magnitude depending on rewarded and unrewarded choices

In our previous study, both M2 and DS ensembles displayed outcome-related activity (Handa *et al*., 2021). As expected, the TCA of the M2-DS ensemble revealed a change in trial factors depending on the outcome (rewarded and unrewarded events) at the single-trial level (Fig. 4A–C). A representative TCA component showcased an increase in within-trial activity (temporal factor) following the response (Fig. 4B). The trial factor highly deviated in unrewarded choice trials without a bias of laterality (Fig. 4C and D) (Left choice: one-way ANOVA: *P* <10^-18^, *post hoc* Dunnett test, Rep-1 & rwd vs. Rep & unrwd: *P* <0.0001, Rep-2 & rwd vs. Rep & unrwd: *P* <0.0001, Rep-3 & rwd vs. Rep & unrwd: *P* <0.0001; Right choice: one-way ANOVA: *P* <10^-27^, *post hoc* Dunnett test, Rep-1 & rwd vs. Rep & unrwd: *P* <0.0001, Rep-2 & rwd vs. Rep & unrwd: *P* <0.0001, Rep-3 & rwd vs. Rep & unrwd: *P* <0.0001). Outcome-selective TCA components were frequently observed in both left- and right-choice trials (Fig. 4E), in contrast to the choice pattern-selective TCA component discussed earlier. The population of Z-scores for the outcome-selective TCA trial factor was significantly higher in the repetitive but unrewarded-choice trials than in the repetitive and rewarded-choice trials (Fig. 4F). This finding suggests that the M2-DS ensembles differentially encoded outcome information in a given trial.

**Figure 4.**
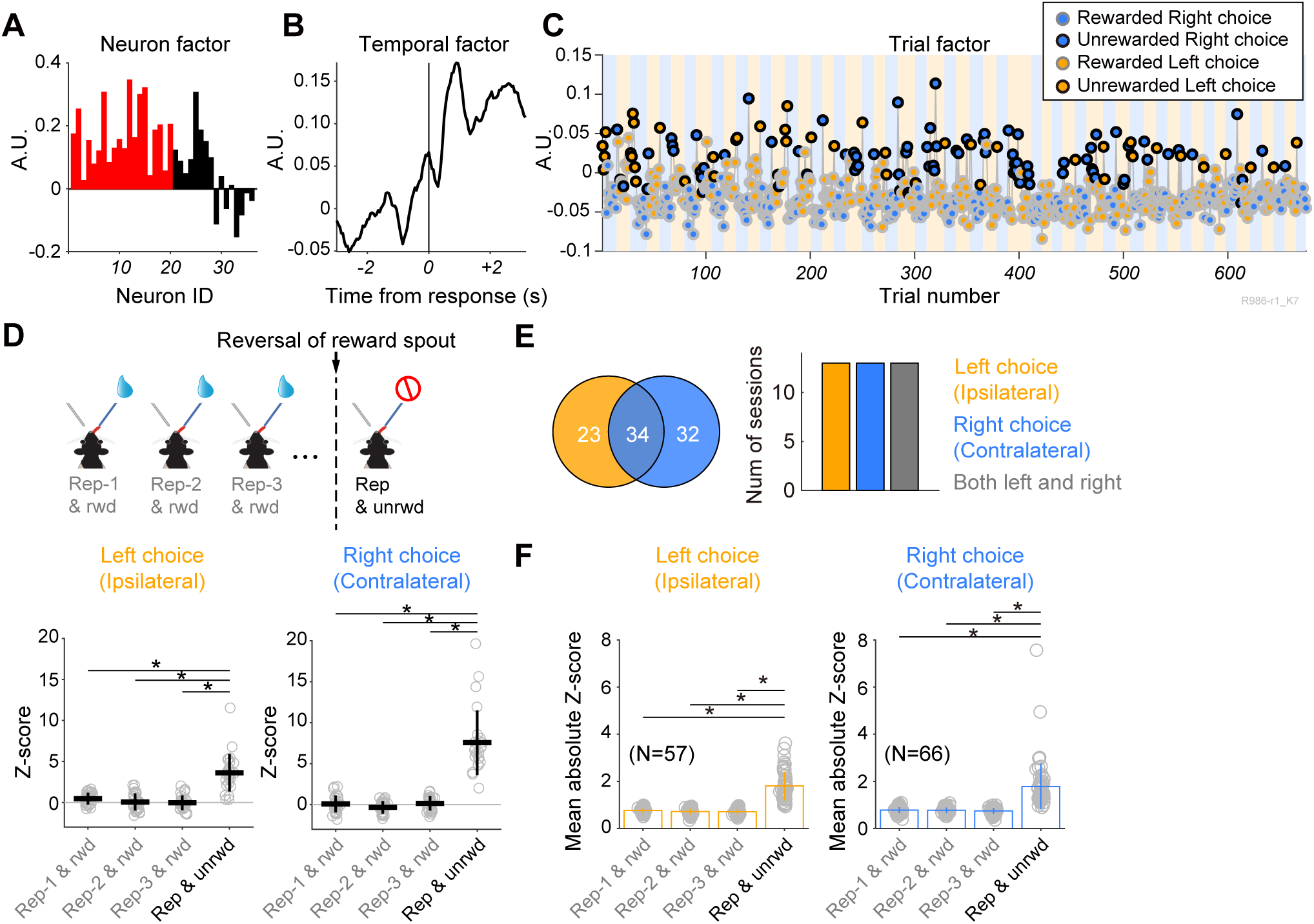
Trial factor of a TCA component distinguishing between rewarded and unrewarded choice trials. **(A)** Neuron factor, **(B)** temporal factor, and **(C)** the trial factor of the TCA component, revealing substantial variance during unrewarded choice trials. In the trial factor, different colours represent choice patterns, with edge and face colours denoting outcomes (rewarded: grey, unrewarded: black) and choice positions (orange: left, blue: right), respectively. **(D)** Top: A schematic illustration of outcome conditions used for statistical estimation of differences in trial factors of repetitive choice trial between rewarded and unrewarded outcomes. A vertical dash line indicates the reversal of reward position from the right spout to the left spout. The trial sequence illustrates a series of repetitive choice trials after a switch choice trial with a rewarded outcome (Rep-1 & rwd, Rep-2 & rwd, and Rep-3 & rwd), as well as repetitive choice trials without reward outcomes after the reversal of the reward-position (Rep & unrwd). Bottom: Z-scores of trial factors computed in Rep-1 & rwd, Rep-2 & rwd, and Rep-3 & rwd, and Rep & unrwd are shown in the session depicted in panel C when the animal made left and right choices. The statistical significance of the difference is assessed through one-way ANOVA followed by *post hoc* Dunnett test (*: *P* <0.05). **(E)** The Venn diagram displays the number of components revealing significantly different trial factors between rewarded and unrewarded repetitive choice trials at left choice (orange), right choice (blue), and both (merge). The bar graph indicates the number of sessions where components showed significant differences between rewarded and unrewarded repetitive choice trials. **(F)** Population data of TCA components revealing a significant difference in Z-scores between rewarded and unrewarded repetitive choice trials. Individual dots represent the mean of absolute Z-score per TCA component, with bar graphs depicting the mean and SD. Statistical significance was assessed through one-way ANOVA followed by *post hoc* Dunnett test (*: *P* <0.05).

### Differential within-trial activity in choice-pattern selective and outcome-selective TCA components

The functionally distinct TCA components likely reflect characteristics revealing different roles of the M2-DS ensemble in adaptive outcome-based decision-making. To address this question, we examined the differences in within-trial activity (temporal factor) between the choice-pattern and outcome-selective TCA components. Most choice-pattern-selective TCA temporal factors were activated before the response and exhibited decreased activity after the response (Fig. 5A). Conversely, >50% of the outcome-selective TCA temporal factors showed an increase in ensemble activity following the response (Fig. 5B). On average, these functionally distinct TCA components displayed opposite trends in temporal factors before and after the response (Fig. 5C). We quantified these temporal differences between choice-pattern-selective and outcome-selective TCA components by comparing the peak time of the temporal factor. The peak time of the temporal factor of choice-pattern selective TCA components (mean ±SD: left choice trials, -0.473 ±1.78 s, right choice trials, - 0.608 ±1.51 s) was significantly earlier than that of outcome-selective TCA components (left choice trials, 0.696 ±1.61 s, right choice trials, 0.525 ±1.51 s) (Mann-Whitney U-test, left choice trials: *P* = 0.0189, right choice trials: *P* = 0.00401). This finding suggests that the M2-DS ensemble plays a temporally distinct role in adaptive choice behaviour by detecting outcomes after the choice response and flexibly making action decisions (such as repeating or switching choice) based on the outcome information.

**Figure 5.**
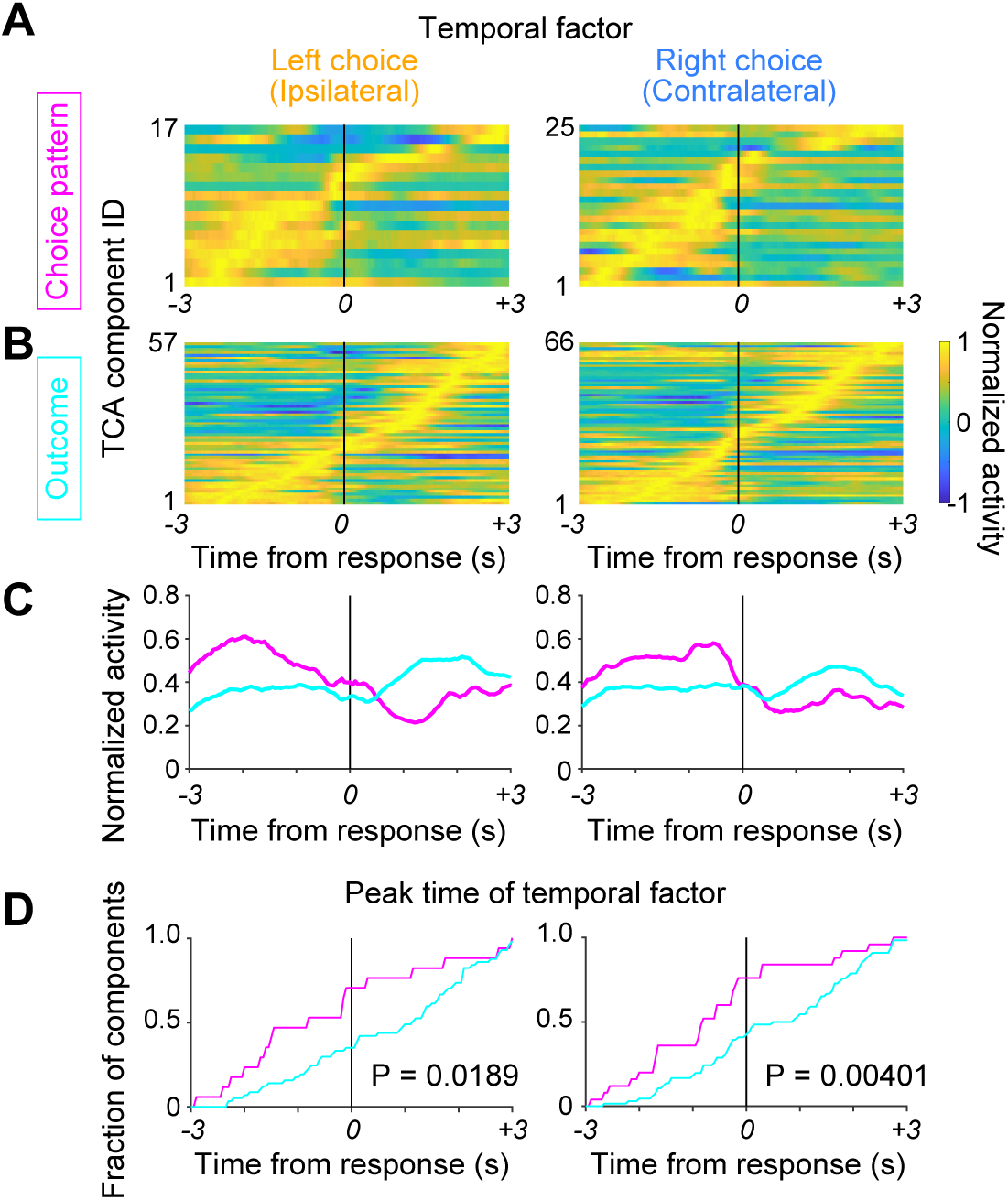
Differential within-trial activity of M2-DS ensembles between choice-patten and outcome selective TCA components. **(A)** Collective normalized temporal factors (within-trial activity) of TCA components with a significant difference in Z-scores of trial factor between repetitive and switch choice trials at left (ipsilateral) and right (contralateral) choice trials. **(B)** Collective normalized temporal factors of TCA components exhibit a significant difference in Z-scores of trial factors between rewarded and unrewarded repetitive choice trials at left (ipsilateral) and right (contralateral) choice trials. **(C)** Averaged normalized temporal factors related to choice patterns (magenta) and outcomes (cyan). **(D)** Cumulative summation curve of peak time of temporal factors of choice-pattern selective TCA components (magenta) and outcome-selective TCA components (cyan). Statistical significance was assessed using the Mann-Whitney U-test.

### Comparison of TCA components based on M2- and DS-alone ensembles with TCA components based on M2-DS ensemble

If the features of ensemble activity of M2 and DS neurons were heterogeneous, or spike activity was not well coordinated between M2 and DS, the application of TCA to the concatenated ensemble activity of M2 and DS neurons (the M2-DS ensemble) could potentially attenuate task-related signals compared with TCA on ensemble activity in each region separately (Runyan *et al*., 2017). To explore this possibility, we investigated whether the choice-pattern- and outcome-selective TCA components derived from the M2-DS ensemble provided worse task-related signals than those obtained from ensembles in M2 or DS alone. TCA was separately applied to the M2 and DS ensembles over the same data used for the M2-DS ensemble analysis. Choice-pattern- and outcome-selective TCA components were observed for both M2 and DS ensembles. Regarding choice-pattern-selective TCA components, in left (ipsilateral) choice trials, the total number of detected TCA components was slightly fewer in the M2-alone ensemble (n=4), whereas that of the DS-alone ensemble (n=15) was comparable to that of the M2-DS ensemble (n=17). In right (contralateral) choice trials, the M2-DS ensemble yielded more choice-pattern-selective TCA components (n=25) than the M2 (n=15) and DS (n=14) ensembles alone (Fig. 6A). We compared the absolute Z-scores of the trial factors in switch choice trials among the M2-DS, M2, and DS ensembles. These values showed similarity among the M2-DS, M2-alone, and DS-alone groups in both left (one-way ANOVA, *P* = 0.665) and right choices (one-way ANOVA, *P* = 0.0587) (Fig. 6B). Regarding outcome-selective TCA components, the total number of detected TCA components (left/right choice: n=58/67) based on the M2-DS ensemble was slightly higher than that based on the M2-alone (left/right choice: n=43/55) and DS-alone (left/right choice: n=40/39) ensembles (Fig. 6C). The Z-scores of the trial factor at unrewarded repetitive choice were also similar among the three datasets in left-choice trials (one-way ANOVA, *P* = 0.184) and right-choice trials (one-way ANOVA, *P* = 0.327) (Fig. 6D). This finding indicates that TCA using the combined M2 and DS ensembles did not attenuate task-related signals; instead, it may not only reflect the features of either the M2 or DS ensemble but also the features of the combined M2 and DS ensemble activity.

**Figure 6.**
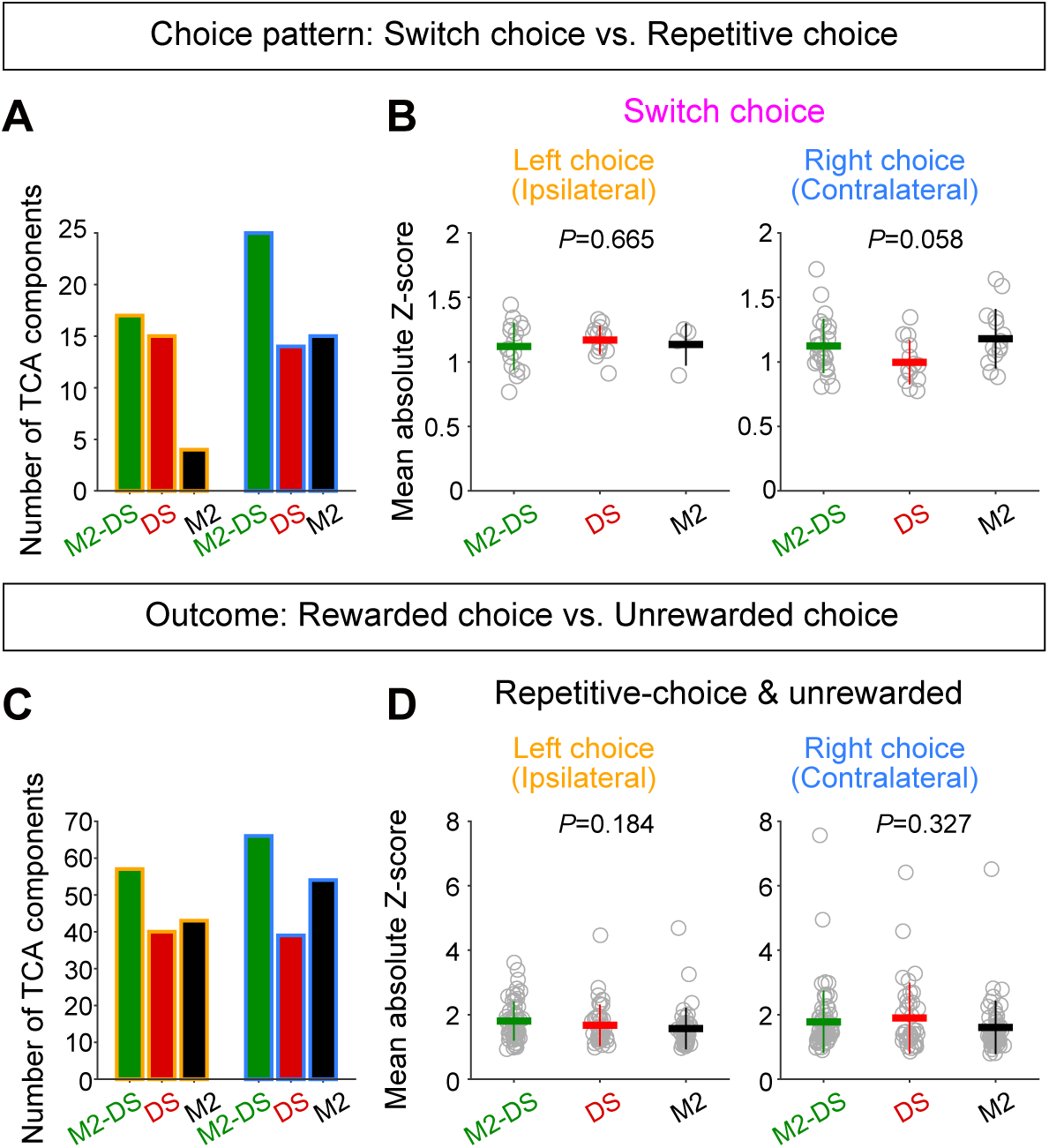
Comparison of TCA component based on M2-DS ensembles with those based on M2- or DS-alone ensemble. **(A)** The total number of TCA components revealing significant differences between repetitive and switch choice trials is shown for M2-DS ensembles (green), M2-alone ensemble (black), and DS-alone ensemble (red) across 14 sessions. **(B)** A comparison of mean absolute Z-scores of trial factors at switch trials for TCA components revealing significant differences between repetitive and switch choice trials among M2-DS ensembles (green), M2-alone ensemble (black), and DS-alone ensemble (red). **(C)** The total number of TCA components revealing significant differences between rewarded and unrewarded repetitive choices is displayed for M2-DS ensembles (green), M2-alone ensemble (black), and DS-alone ensemble (red). **(D)** A comparison of mean absolute Z-scores of trial factors at unrewarded repetitive choice trials for TCA components revealing significant differences between rewarded and unrewarded repetitive choices among M2-DS ensembles, M2 ensemble alone, and DS ensemble alone. Horizontal and vertical lines represent mean and SD, respectively.

## Discussion

In this study, we demonstrated distinct representations of decomposed M2-DS ensemble activity for various task-relevant behaviours at the trial level in rats performing an outcome-based choice task using the TCA approach. Although TCA is an unsupervised method, in contrast to linear discriminant analysis and linear regression, it automatically identifies choice-position, choice-pattern, and outcome-specific patterns. Choice-position-selective TCA components (the choice-position-specific spatiotemporal neural dynamics) revealed dynamic changes in selectivity over trials, which were correlated with changes in behavioural variables, such as RTs, and/or the number of licks after the response. The TCA revealed neural dynamics showing selectivity for choice patterns (repeat and switch choices), even when the choice position and outcome were identical. Choice pattern- and outcome-selective neural dynamics revealed functionally distinct within-trial activity. Choice-pattern selective within-trial activity increased activity earlier (before the choice response), whereas outcome-selective within-trial activity increased activity later (after the choice response). TCA application on M2-DS ensemble activity tended to yield more task-related neural dynamics than TCA application on M2 or DS alone.

The rodent M2 integrates sensory and outcome information to make decisions regarding motor planning (Barthas & Kwan, 2017). The activity of M2 neurons represents laterality in the body (Sul *et al*., 2011; Erlich *et al*., 2011), forelimb movements (Soma *et al*., 2017), and tongue movements (Siniscalchi *et al*., 2016; Handa *et al*., 2017, 2021; Kurikawa *et al*., 2018). Additionally, the M2 is implicated in outcome evaluation for action (Sul *et al*., 2011; Gremel & Costa, 2013; Handa *et al*., 2021) and in the neuronal encoding of outcomes, such as reward and non-reward events (Sul *et al*., 2011; Handa *et al*., 2021). The rodent DS serves a significant gateway of the basal ganglia, receiving major synaptic inputs from the cerebral cortex (Kincaid *et al*., 1998; Reep *et al*., 2003; Cheatwood *et al*., 2003; Wall *et al*., 2013; Hintiryan *et al*., 2016). Neuronal activity in DS similarly shows the laterality of the choice response (Kim *et al*., 2009; Handa *et al*., 2021) and the value of the outcome (Kim *et al*., 2009; Nonomura *et al*., 2018). Therefore, similar behaviour-related signals are observed in the two regions. To gain a deeper understanding of the mechanisms underlying cross-region transmission between M2 and DS in outcome-based decision-making, we used Fisher’s linear discriminant (FLD) for dimensionality reduction of ensemble activity. In our previously work, we extracted the dynamic characteristics of the ensemble activity in each region and compared these characteristics between M2 and DS. Neural trajectories in the M2 and DS revealed temporally similar task-related signals, such as choice position and outcome. Precise spike synchrony between M2 and DS neurons emerges more frequently when task performance is superior (Handa *et al*., 2021). Consistent with our previous data, a recent study demonstrated that cortical representation is topographically reflected in the striatal subregion (Peters *et al*., 2021). Our current findings demonstrate that the M2-DS ensemble activity can be deconstructed into distinct functions— specifically, choice position, choice pattern, and outcome—without compromising the representation of decision-related signals in each region. This evidence suggests that the ensemble activity of interconnected regions (the M2-DS ensemble) effectively processes similar decision-related signals in a cooperative manner for further processing through downstream structures in the basal ganglia for motor selection.

Although our previous work successfully identified temporally parallel processing of information related to choice position and outcome between the M2 and DS using FLD, we encountered challenges in identifying choice-pattern selective activity at the population activity level (Handa *et al*., 2021). Similar to conventional principal component analysis, FLD utilizes trial- averaged data to compute a vector of hyperplanes, maximising the degree of discrimination between two conditions (left and right choices) (Bishop, 2006). The assumption of trial averaging is that trial-by-trial variability is task-irrelevant noise. Conversely, TCA considers trial-by-trial variability to extract features of population activity at a single-trial level in an unsupervised manner (Williams *et al*., 2018). Our current results not only demonstrated choice-position and outcome-selective M2-DS ensemble activity but also revealed choice-pattern-selective activity based on trial-by-trial analysis. These choice-pattern selective neural dynamics were observed even in neural dynamics based on M2 or DS ensemble activity alone, suggesting that the choice pattern is commonly processed in both M2 and DS. Choice-pattern-selective neural dynamics of M2-DS ensemble activity were found more frequently than those of M2 or DS alone in contralateral choice trials. In this case, dimensionality reduction in an unsupervised manner enables the uncovering of the latent features of the neural ensemble. This intriguing component of the M2-DS ensemble activity may reflect cognitive features for action selection based on outcome rather than motor commands as the incoming choice position is identical, but the difference lies in selecting the spout position by deciding whether to repeat the previous choice or switch from the preceding choice in the outcome-based choice task. In support of this interpretation, previous studies have indicated that M2 is involved in cognitive switching between sensory-guided and automated choice rules (Siniscalchi *et al*., 2016), as well as flexible visual categorisation (Wang *et al*., 2020). The DS is additionally implicated in the processing of action selection based on outcome probability or value (Nonomura *et al*., 2018; Cox & Witten, 2019). Our findings are consistent with these neuronal functions in both the M2 and DS, extending to M2-DS ensembles.

Choice-position-selective TCA neural dynamics revealed trial-to-trial fluctuations in choice-position selectivity, which correlated with variable behavioural parameters. The correlation with the number of licks may indicate the significance of the M2-DS ensemble activity in relation to changes in motivation to participate in the task, given the decrease in the number of licks observed in the later period of the session. This newly observed result is also attributed to the trial-based TCA approach.

A recent study demonstrated that synchronous spike activations across regions, including the medial prefrontal cortex and the downstream dorsomedial and ventral striatum, emerge with behavioural correlations contingent on task demands during the T-maze task. Different firing assemblies were observed at various times within the task (Oberto *et al*., 2022). In our study, the within-trial activity revealed distinct characteristics, such as choice-pattern-selective neural dynamics and outcome-selective neural dynamics, which varied temporally. This suggests that the M2-DS ensemble serves different functions at different time points. Choice-pattern-selective neural dynamics may be modulated before the choice response, influencing the decision to repeat or switch, whereas the outcome-selective component could activate after the response to monitor the outcome. In essence, the M2-DS ensemble contributes to temporally distinct roles in adaptive choice behaviour by detecting outcomes after choice response and flexibly making decisions of action (repeating or switching choices) based on the outcome information. Growing evidence indicates that frontal cortex-striatal ensemble coactivation emerges at specific times and plays specific roles in ongoing goal-directed behaviour.

## Acknowledgments

We express our gratitude to R. Harukuni for her invaluable support in behavioural training and histological analysis

